# Design Tradeoffs in High Resolution, Large Field-of-View 3D All-Optical Control of Neural Circuits

**DOI:** 10.64898/2026.10.28.762619

**Authors:** Barbara Gruszka, Masashi Hasegawa, Ian Antón Oldenburg

## Abstract

Multiphoton holographic optogenetics is an emerging technology that offers the ability to probe neural circuits with unprecedented temporal and spatial precision. When combined with *in vivo* imaging, it enables a host of experiments that were not previously possible. While this approach grows in popularity, the ‘one-size-fits-all’ approach of most microscopes leads to systems that are not optimized for many experimental paradigms. Focusing on scanless temporally focused holographic optogenetic approaches, we explore the tradeoffs between resolution and field of view in different microscope designs. We show that simple changes in optical elements and lens order can create systems with both improved accessible range and resolution. Taking advantage of newer, high NA, 2P-optimized, low-magnification objectives, we demonstrate cellular-resolution holographic control over neurons separated by more than 1.4 mm in a single hologram, significantly expanding the spatial extent accessible for simultaneous neural manipulation in a conventional microscope. These changes are built upon a simple framework that leverages head-to-head system comparisons as a critical approach for evaluating the true advantages and limitations of an optical design, which could be incorporated into a wide variety of microscopes.

## Main

Multiphoton holographic optogenetics is an emerging technique that offers the ability to probe neural circuits, thereby enabling researchers to address previously intractable questions in systems neuroscience. Holographic approaches span several modalities, encompassing work in mouse visual cortex^1–10^, somatosensory cortex^11–14^, olfactory cortex^15^, motor cortex^16–19^, hippocampus^20,21^, as well as in parietal and auditory cortices^22^.

Despite the growing popularity of this technique, most multiphoton stimulation systems take on relatively few configurations, regardless of the specific application. While the advantages of different stimulation techniques has been explored^23–27^, experimental studies exploring how distinct optical paths of a given architecture can vary the results are scarce. As a result, researchers new to holographic optogenetics are often unaware of how to optimize their multiphoton microscopes for their specific application. For instance, the microscope objective and upstream light path design are simple but critical decisions when building a multiphoton holographic system. These decisions are necessary to strike an optimal balance between a larger field-of-view (FOV) *and* resolution.

We focus on a scanless, temporally focused, 3D multiphoton holographic optogenetic approach (3D-SHOT). When combined with fast opsins such as ChroME2s, 3D-SHOT offers exquisite temporal resolution of evoked action potentials^3,28–33^. Furthermore, scanless approaches are not constrained by galvo mirror size, enabling true synchronous stimulation across a large field of view unlike scanning systems, which can compromise the FOV to keep the targets contained on a small mirror.. We identify a key change in the 3D-SHOT system’s stimulation optical path that improves its use with lower magnification objectives (e.g. 10x objectives) for greater spatial resolution and larger field-of-view of holographic stimulation.

While there have been several efforts to compare objectives for multiphoton and Fourier microscopy^34–36^, only a few reports utilize standardized methodology for empirically measured optical point-spread functions or other optical properties^30,37,38^. Moreover, multiphoton optogenetic stimulation involves a special use case, where the tradeoffs of resolution, field of view, and power throughput may not have the same requirements as in imaging only modalities. Furthermore, while mathematical modeling of objectives and light path changes provides a starting point, empirical validation and in-vivo testing remain indispensable.

By modifying the standard 3D-SHOT light path and using a low magnification objective we retain cellular resolution while expanding the holographically accessible FOV across over 1.4mm of tissue. This change offers an increase in accessible area of over 40% from previous state of the art^6^. While most multiphoton holographic photostimulation studies utilize a field of view with a diameter ≤1mm, several groups expand this with asynchronous stimulation approaches either by raster scanning^39^, or by redirecting a smaller FOV with galvos^40,41^.

Altogether, we seek to relate these changes to the experimental needs of independent researchers and provide a strategic guide for the advancement of multiphoton holography.

## Results

### Calibration and Measurement

A 3D-SHOT multiphoton stimulation path was constructed as previously described^29,31^ (Fig.1A-B). In 3D-SHOT, the size of the photostimulated target is set by the light path, without the need for scanning. Throughout this manuscript, we measure the three-dimensional location and size of photostimulation targets by projecting holograms containing multiple targets onto a thin fluorescent slide conjugated to a substage camera (Fig.1C). We move the targets axially in both directions, with respect to the slide and substage camera, by defocusing with the microscope objective. The slide is imaged with a low magnification (4x) objective to capture holographic targets throughout a large area (Fig. 1D, left). For each target, we measured its evoked fluorescence intensity at several depths to create a three-dimensional representation of the stimulated volume (Fig. 1D, right). We use fluorescence as our measure of two-photon efficacy, extrapolating the peak intensity and full width at half maximum (FWHM), i.e. an optical point spread function (OPSF), to quantify the radial and axial resolution of each target (Fig. 1E).

**Figure 1.**
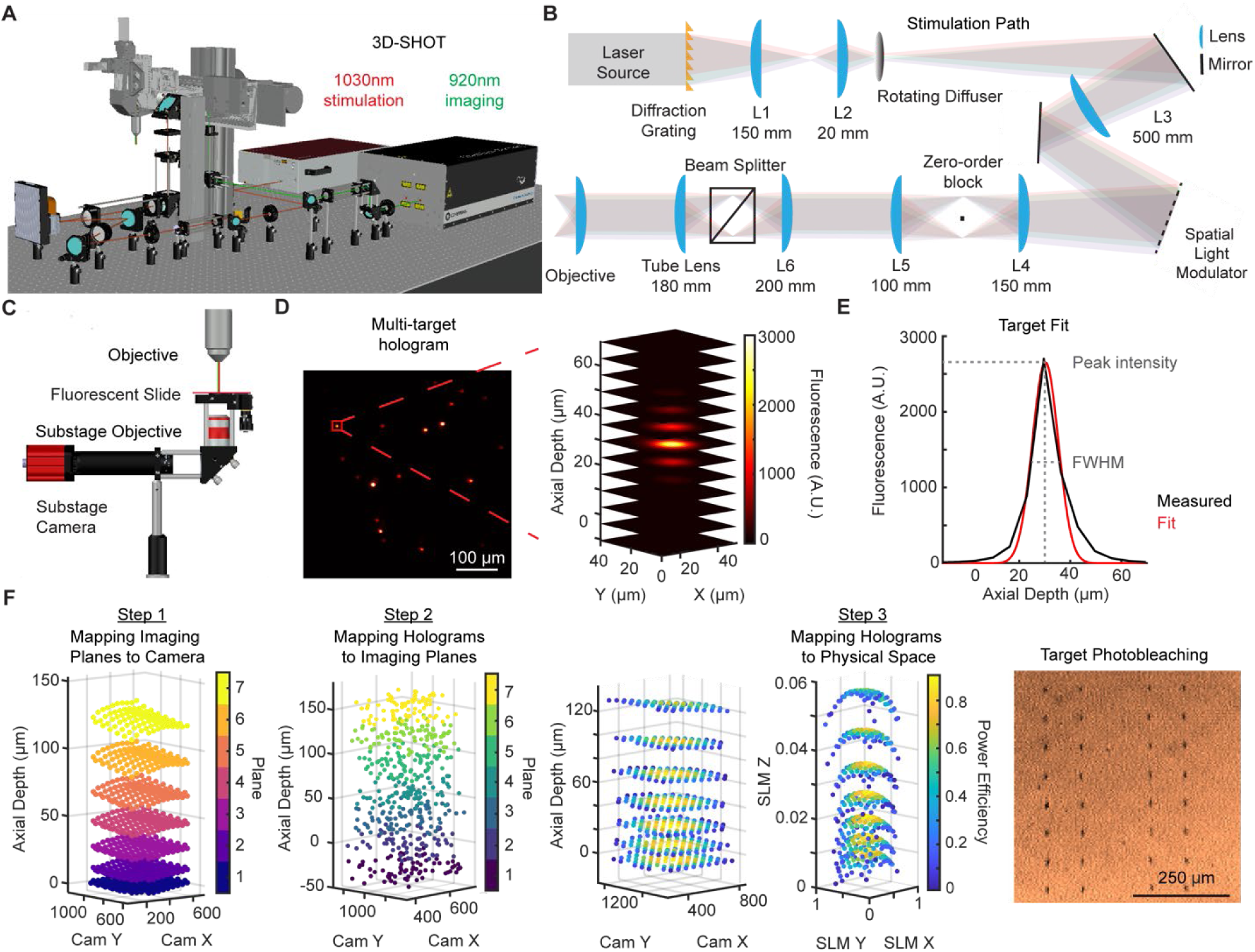
**A)** Modeled multiphoton 3D-SHOT microscope system, with stimulation (red) and imaging (green) optical pathways originating at the 1030nm photostimulation and 920nm imaging lasers. **B)** Schematic of stimulation path. **C)** calibration set-up, consisting of the microscope objective and a thin, fluorescent slide atop a second objective conjugated to a substage camera. **D)** Left, substage camera view of a multi-target hologram on a fluorescent slide. Right, a single target from the hologram visualized across axial depth. **E)** example fluorescent measurements across axial depth (‘Measured’, black), fitted to a gaussian (‘Fit’, red), and full-width-half-max calculation of the optical point-spread function, ‘OPSF’. **F)** Step 1: Detected location of 7 imaging planes, by the substage camera. Step 2: 700 targets spanning the field of view of the imaging planes are divided into groups of 20, by estimated depth. Each target is independently detected by the substage camera and its location, intensity, and size are recorded. Step 3: Holograms targeting the center of each of the 7 planes are generated and mapped to ‘SLM space’. High intensity illumination of each target creates a bleached spot in the fluorescent slide, that can be detected and registered via 2P-imaging (‘Target Photobleaching’).

Holographic targets were mapped to physical coordinates using a three-step calibration procedure (Fig.1F, Methods) adapted from those described previously^7,30^. This pipeline registers individual imaging planes to the substage camera (Step 1), maps target location, intensity, and resolution across the 3D volume to build a polynomial model relating ‘SLM coordinates’ to camera space (Step 2), and uses bleached targets to establish a second model relating ‘SLM coordinates’ to physical space on 2p images (Step 3).

### Objective Comparison

We compare two water-dipping objectives: the 20X Olympus (Olympus) and 16X Nikon (Nikon) and two 10X air objectives: the Super Apochromatic (Thorlabs) and Cousa (Pacific Optica). Their reported optical properties are outlined in Supplementary Table 1. Following the alignment of the imaging and stimulation paths, we acquired radial, axial, and power efficiency measurements of targets spanning a depth of -25 to 120 µm for the four objectives. Using the conventional 3D-SHOT light path and standard SLM (HSP1k, Meadowlark Optics, Fig. 1B), the average radial OPSF FWHM was 9.36 ± 0.04 µm (‘Olympus – 20x’), 10.0 ± 0.07 µm (‘Nikon – 16x’), 12.1 ± 0.08 µm (‘ThorLabs – 10x’), and 13.8 ± 0.14 µm (‘Cousa – 10x’) (mean ± SEM; Fig. 2A-D).

**Figure 2.**
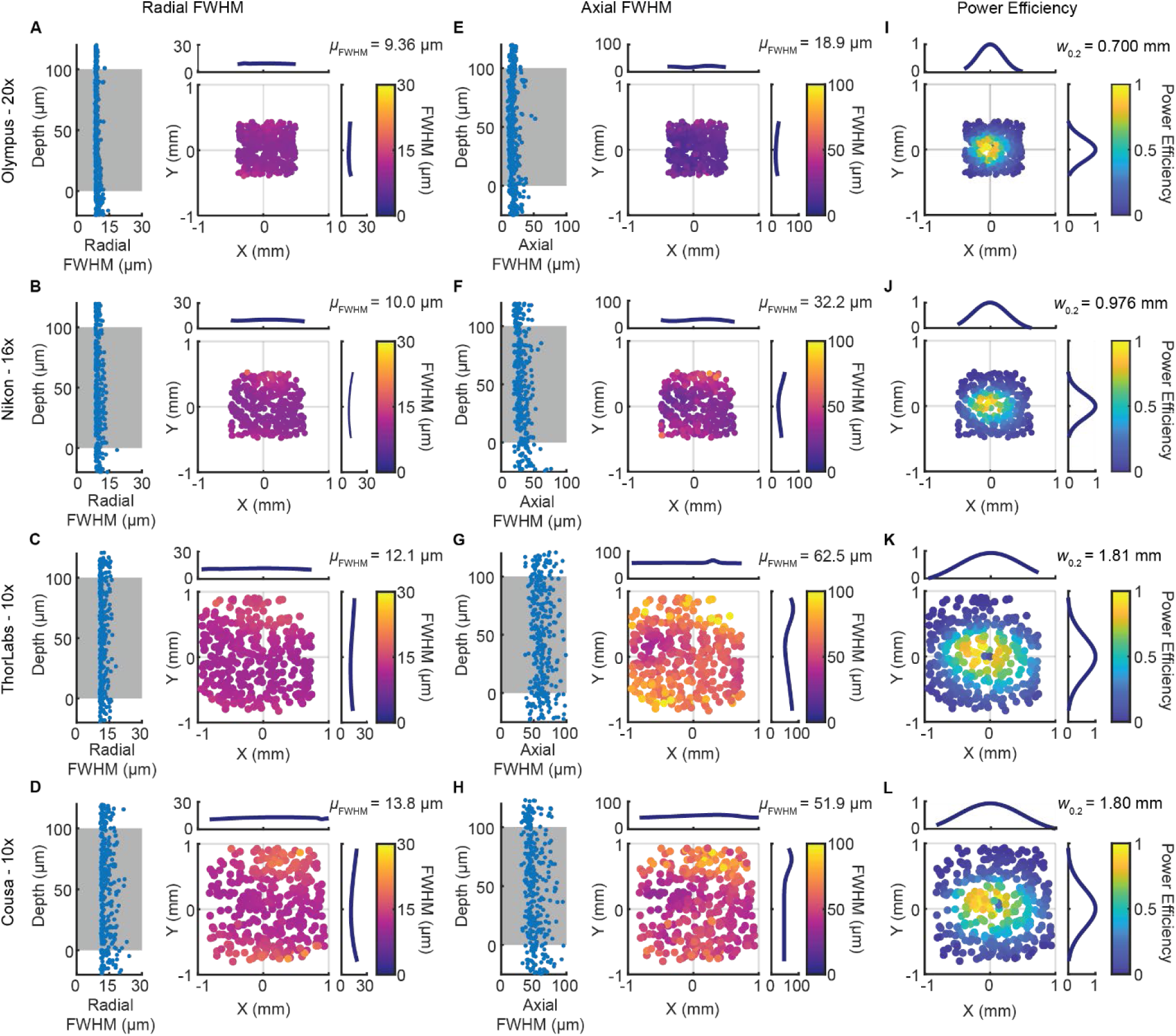
Characterization of FWHM and power efficiency of holographic targets across objective lenses. Each row corresponds to a single objective under the default optical path configuration. **A-D)** Left, radial FWHM profiles (µm) as a function of depth. The gray shaded region indicates the range of axial depths typically utilized in experiments, though the system can accommodate a larger span. Right, radial FWHM profiles across the FOV, with gaussian fits shown in the top (X) and right (Y) panels. µ_FWHM_ indicates the average FWHM measurement. **E-H)** as in A-D but for axial FWHM. **I-L)** Power efficiency across the FOV, with gaussian fits shown in the top and right panels. W_0.2_ denotes the full width at 20% peak power efficiency.

We derived an analytical model to calculate the size of the stimulation beam after changes in relay lenses magnification (see Supplementary Note 1 for a worksheet incorporating this analytical calculation). These empirical measurements agree closely with our analytical model (Methods, Supplementary Table 2). 3D-SHOT uses relay lenses to intentionally create a supra-diffraction-limited stimulation target to illuminate an entire neuron’s soma during an *in vivo* experiment. The analytical model’s target size assumes zero aberration, which represents conditions closer to the center of the FOV at the true focal plane. To better compare experimental and modeled results, we also averaged FWHM values for points with power efficiencies above 20% (Supplementary Table 2).

**T**he average axial OPSF FWHM was 18.9 ± 0.28 µm (‘Olympus – 20x’), 32.2 ± 0.48 µm (‘Nikon – 16x’), 62.5 ± 0.69 µm (‘ThorLabs – 10x’), and 51.9 ± 0.57 µm (‘Cousa – 10x’) (mean ± SEM; Fig. 2E-H). As expected, lower magnification objectives had higher averaged measured axial FWHMs.

As expected, holograms in the center of the FOV exhibit the greatest power efficiency while the edges see a diminished efficiency due to the SLMs deflecting light at a severe angle (Fig. 2I-L). We measure the area that we can effectively target as locations where holograms are at least 20% of the maximum observable intensity. The area of effective control varies by objective: 0.70 mm (‘Olympus – 20x’), 0.98 mm (‘Nikon – 16x’), 1.81 mm (‘ThorLabs – 10x’), and 1.80 mm (‘Cousa – 10x’) diameter (Fig. 2I-L, W_0.2_).

To evaluate the influence of the off-axis distance and depth on the holographic targets, data were analyzed using a cross-objective linear mixed model (SFig. 1). FOV zones were defined as concentric 2D annular rings extending outward from the center in 200 µm radial increments. For all targets with power efficiencies ≥ 20%, spatial resolution exhibited relative consistency across FOV zones and depth. Pooled radial FWHM showed no significant dependence on FOV zone (β = +0.118 µm/mm, p = 0.584; SFig. 1A-D) and a very minor axial depth sensitivity (β = -2.07 µm/mm, p = 0.0044). Similarly, axial FWHM showed no significant dependence on FOV zone (β = -2.97 µm/mm, p = 0.112; SFig. 1E-H), alongside a slight axial depth variation (β = -29.0 µm/mm, p = 4.68e-6). Conversely, power efficiency roll-off was overwhelmingly driven by FOV zone (β = - 1.26 mm^−1^, p = 1.63e-139; SFig. 1I-L). Axial depth yielded no statistically significant contribution to power loss (β = +0.031 mm^−1^, p = 0.813). Overall, the holographically addressable volumes are primarily bound by the off-axis power efficiency fall-off rather than depth.

Additionally, we measured the power of each objective in this study in a range from 750 to 1320nm. At 1035nm, the most common wavelength for multiphoton photostimulation, the objectives transmission efficiency ranged between 57 and 94% (SFig. 2).

### *In vivo* demonstration of large field of view stimulation

Our measurements indicate that when combined with a 10x objective, we should be able to activate neurons across a very large area in a single simultaneous hologram. We prepared mice with GCaMP6s and ChroME2s^33^ opsin in excitatory neurons (see Methods), and each opsin-expressing neuron was tested to determine the minimal power necessary to evoke a response, as described previously^7,19,28,29^. To demonstrate the largest area accessible in a single hologram we prepared a 90-target hologram spanning approximately 1.4 mm of the mouse motor cortex. Reliable neural activity was evoked in cells on either side of this hologram; moreover, >80% of targeted neurons responded reliably consistent with previous results^7,19,28,33^.

### Spatial Light Modulator Comparison

As accessible area is a function of the density of pixels on the SLM^42,43^, we tested if we could further increase our FOV by replacing our typical SLM (Meadowlark HSP1k 1024 × 1024 pixels) with one with a higher resolution (Meadowlark HSP1920HB 1920 x 1152 pixels).

We repeated our calibration and measurement process with both SLMs (Fig. 4A-J). We find the average radial and axial OPSF FWHM of the ‘Olympus – 20x’ with the denser HSP1920 SLM to be similar to the standard HSP1k (radial FWHM 9.22 ± 0.08 µm HSP1920, vs 9.36 ± 0.04 µm HSP1k, p = 2.4e-07; axial FWHM 19.9 ± 0.39 µm HSP1920 vs 18.9 ± 0.28 HSP1k, p = 0.06, Fig. 4A, E). The ‘Cousa – 10x’ objective exhibited a modest improvement in resolution with the denser SLM (radial FWHM 10.7 ± 0.08 µm HSP1920, vs 13.8 ± 0.14 µm HSP1k, p = 5.1e-62; axial FWHM 49.8 ± 0.68 µm HSP1920 vs 51.9 ± 0.57 HSP1k, p = 0.0037. Fig. 4B, F).

**Figure 3.**
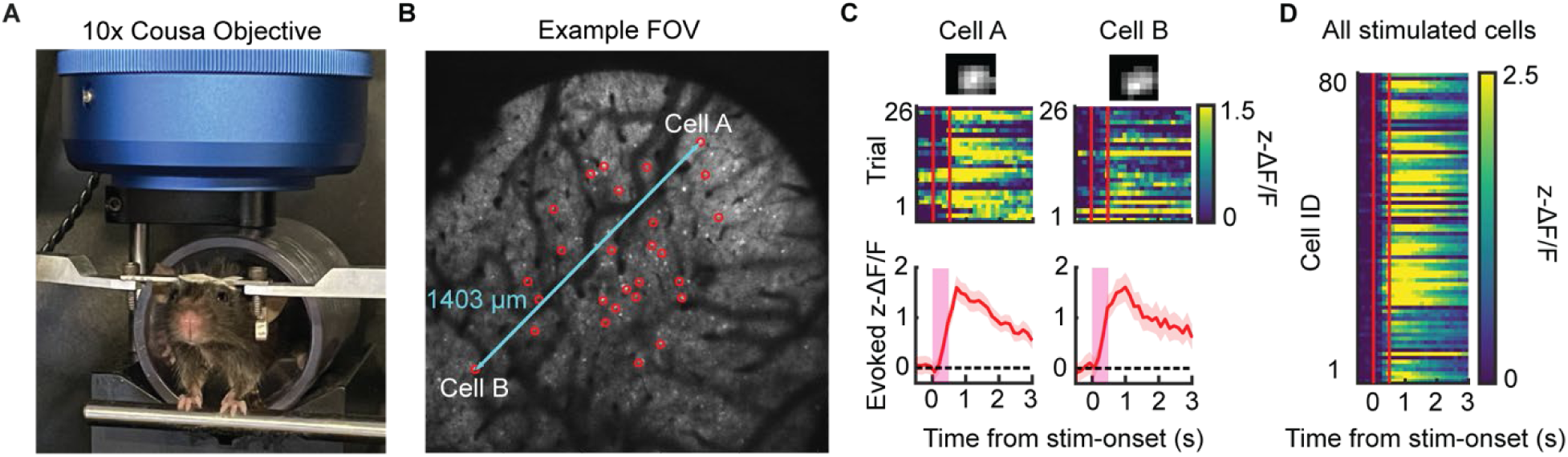
Large FOV stimulation using the 10x Cousa objective. **A)** Image of head-fixed mouse underneath the objective. **B)** Example 10x FOV plane. Red circles indicate stimulated cells in the plane. Light blue line marks the furthest distance (1403µm) between two, individually stimulated cells. **C)** Top: trial-wise, evoked z-ΔF/F calcium activity of example cells A and B. Bottom: trial-averaged activity. **D)** trial-averaged activity of 80 recovered cells out of 90 stimulated cells in the FOV. Representative data from one of two biological replicates averaging to a range of 1.36 mm (N = 2 FOVs, 2 mice).

**Figure 4.**
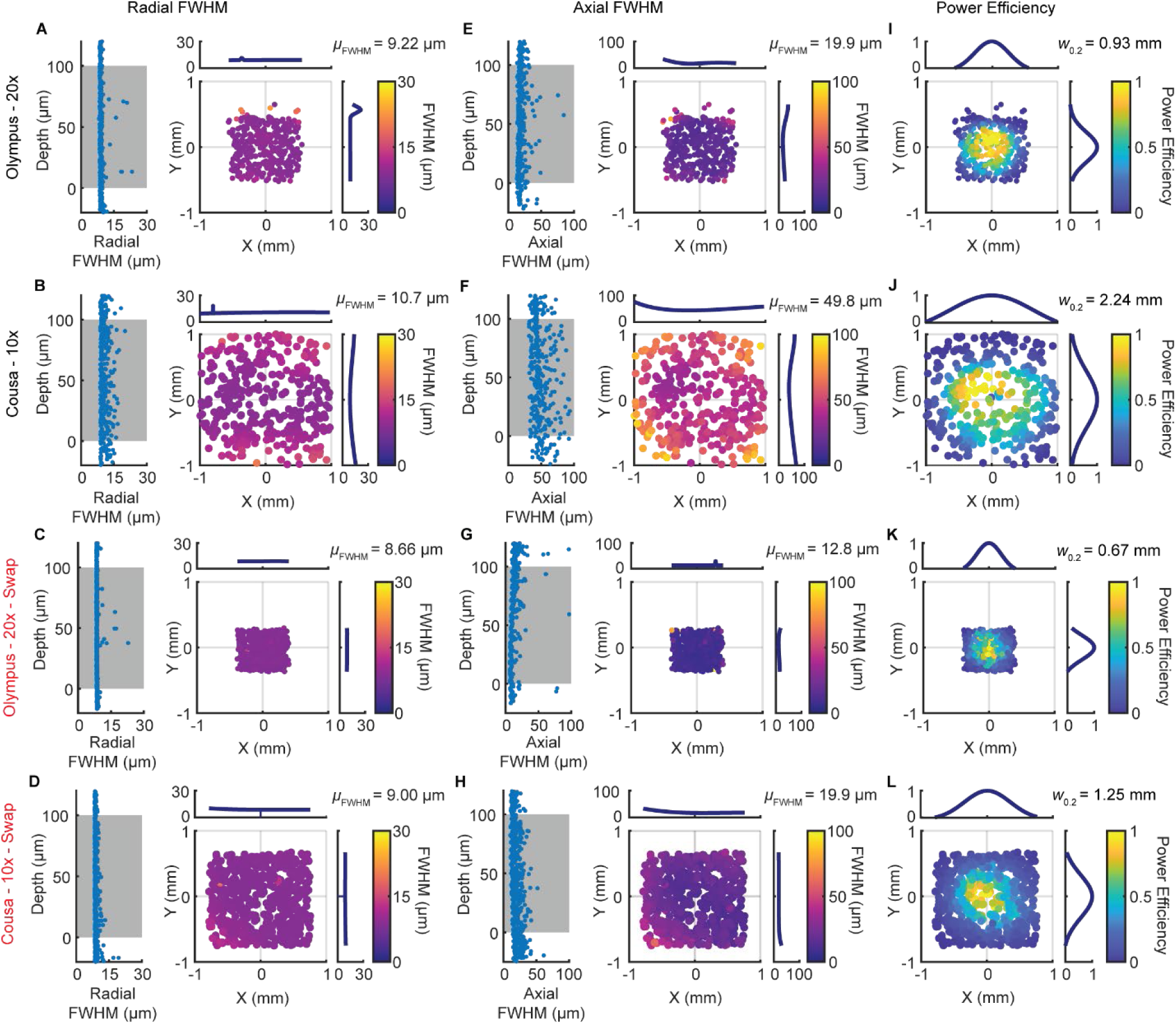
Characterization of FWHM profiles and power efficiency in default and lens-swap (red) conditions for the 20x and 10x objectives using the HSP1920 SLM. Each row corresponds to a single objective-optical path configuration. **A-D)** Left, radial FWHM profiles (µm) as a function of depth. The gray shaded region indicates the range of axial depths typically utilized in experiments, though the system can accommodate a larger span. Right, radial FWHM profiles across the FOV, with orthogonal fits shown in the top (X) and right (Y) panels. **E-H)** Left, axial FWHM profiles across depth. Right, axial FWHM profiles across the FOV, with orthogonal fits shown in the top and right panels. **I-L)** Power efficiency across the FOV, with gaussian fits shown in the top and right panels. *W_0.2_* denotes the full width at 20% peak power efficiency. Up to 700 targets in 20 target hologram configurations illuminate the slide at varying depths.

As predicted, systems with both objectives saw an increased accessible field of view with the denser SLM (‘Olympus – 20x’ W_0.2_ = 0.93mm HSP1920 vs 0.70mm HSP1k, Fig. 4C; ‘Cousa – 10x’ W_0.2_= 2.24 mm HSP1920 vs 1.80mm HSP1k, Fig. 4J). This expanded optogenetically accessible FOV was larger than the full imaging field of view allowed by our microscope (Sutter MOM with a 20° scan angle: ‘Olympus – 20x’ 0.85 µm, ‘Cousa – 10x’ 1.7mm). This leads to a situation where there are hypothetically accessible targets (by resolution and power efficiency) that are not observable while 2p imaging.

### Optimizations for Large Fields of View

While the lower magnification objectives offer extensive accessible FOV, this comes at the expense of single target resolution. To mitigate the impact of this trade-off, we modified the system by swapping lenses 4 (*f* = 150 mm) and 5 (*f* = 100 mm) after the SLM (Fig.1B, Methods). By modeling this change, we expected to minimize the axial FWHM for any objective by modestly reducing the accessible area.

The average radial FWHM for both objectives was similar, but slightly improved, by this ‘swapped’ lens configuration (‘Olympus – 20x’ swapped 8.66 ± 0.04 µm vs original 9.22 ± 0.08 µm, p = 1.5e-77, Fig. 4C; ‘Cousa – 10x’ swapped 9.00 ± 0.04 µm vs original 10.7 ± 0.08 µm, p = 1.2e-89, Fig. 4D). In contrast, the lens swap approach greatly improved axial resolution, especially with the lower power objective (axial OPSF FWHM ‘Olympus – 20x’ swapped 12.8 ± 0.38 µm vs original 19.9 ± 0.39 µm, p = 2.1e-79, Fig. 4G; ‘Cousa – 10x’ swapped 19.9 ± 0.28 µm vs original 49.8 ± 0.68 µm, p = 1.3e-140, Fig. 4H).

As expected, the swap configuration reduced the accessible FOV (‘Olympus – 20x’ W_0.2_ = 0.76mm, Fig. 4K; ‘Cousa – 10x’ W_0.2_ = 1.25, Fig. 4L). This reduced accessible area is still very similar to the maximal imaging field of view in many microscopes, and larger than most reported holographically accessible FOVs. Thus, we demonstrate that by swapping just two lenses, we can retain single cell resolution with a low magnification objective, over much larger FOVs than typically possible.

### Physiological Point-Spread Function

While the optical point-spread function (OPSF) serves as a reliable benchmark for holographic spot size, it strictly defines the physical limits of the microscope optics and the quality of the alignment^37^. As a result, we focus on the *in vivo* physiological point-spread function (PPSF) which describes the functional resolution of stimulation. Here, ensembles of ten opsin-positive neurons are targeted for stimulation, and off-target activity is quantified as the hologram is systematically offset in both the axial and radial dimensions (Fig. 5A, see Methods). We record calcium activity and stimulate cells in the motor cortex (Fig. 5B), as characterizing the PPSF of targeted stimulation is critical for accurately evaluating its effect on surrounding non-targeted cells.

**Figure 5.**
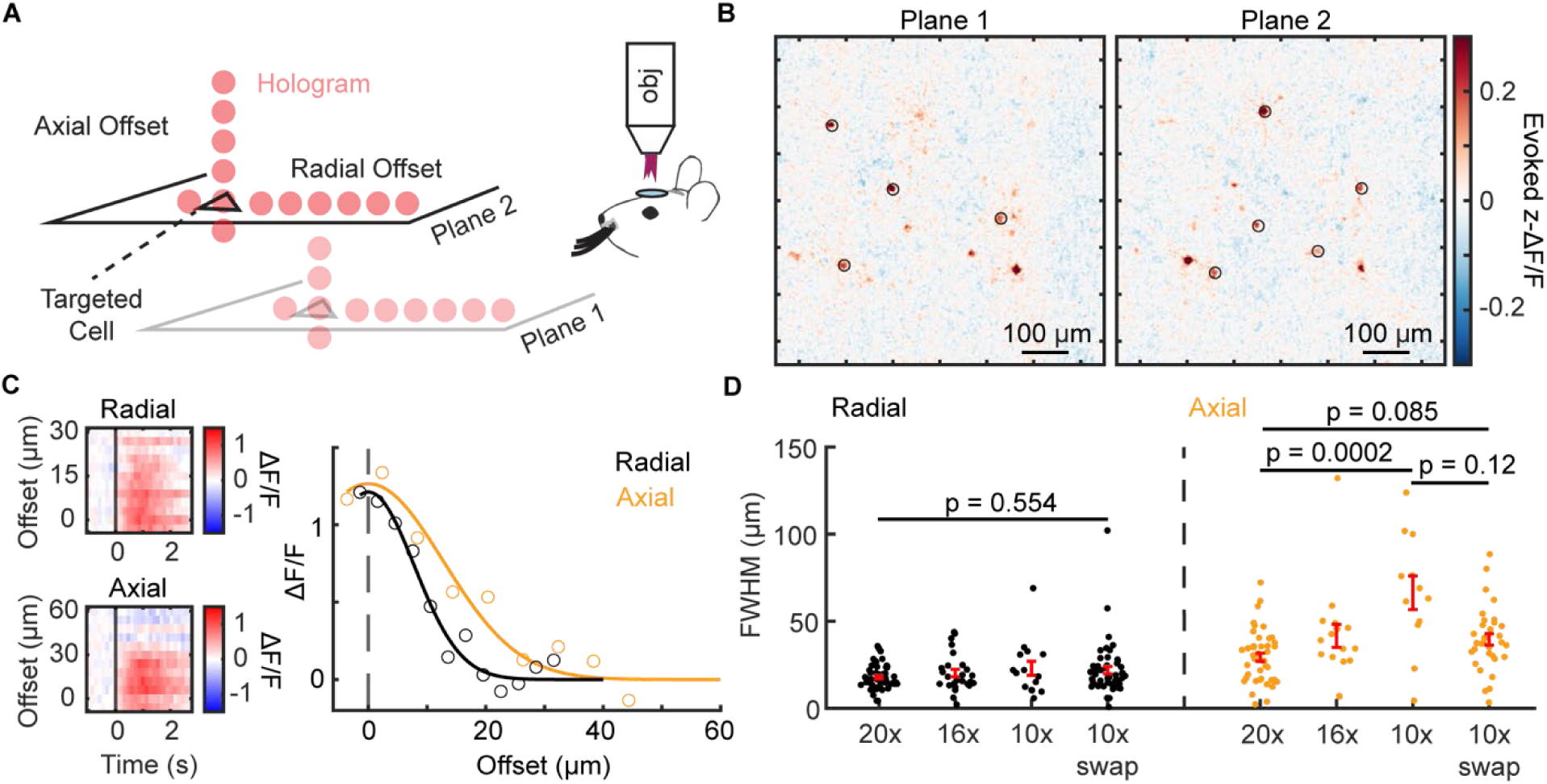
Characterization of the PPSF *in vivo*. **A)** Schematic of the experimental design. Ten cells across two planes are selected for stimulation, followed by stimulation with holographic targets offset in either radial or axial directions. **B)** Example FOV of averaged evoked activity of simultaneously stimulated targets, indicated with black circles. **C)** Left, example trial-averaged activity at each radial (top) and axial (bottom) offset for a single cell. Right, gaussian fits for a single cell, denoted by black and orange lines, respectively. Open circles indicate trial-averaged averaged radial and axial activity at each offset. **D)** Measured radial and axial FWHM using 20x, 16x, and 10x objectives, as well as the 10x objective under the swap condition. Irrespective of magnification, radial FWHM is not significantly different between objectives in either default or swap configurations. In contrast, 20x and 10x objectives are significantly different in axial FWHM. The swap configuration decreases the 10x objective’s axial FWHM to levels comparable to the 20x and 16x objectives.

We measure the PPSF for the 20x, 16x, and 10x objectives in default configuration, and the 10x objective in the swap configuration. For each cell, we fit the evoked z-ΔF/F at each offset step with a gaussian to establish the effective resolution, i.e. the PPSF (Fig. 5C). The 20x, 16x, and 10x objectives yield average radial PPSF FWHMs of 17.9 ± 0.92 µm (‘20x’, N = 55 cells, 4 FOVs, 2 mice), 20.29 ± 2.12 µm (‘16x’, N= 26 cells, 2 FOVs, 2 mice), and 23.0 ± 4.07 µm (‘10x’, N = 15 cells, 2 FOVs, 2 mice) (mean ± SEM; Fig. 5D, radial). Under the 10x swap configuration, the radial PPSF FWHM is also comparable, with an estimate of 21.9 ± 2.09 µm (‘10x-swap’, N = 51 cells, 8 FOVs, 4 mice). The radial PPSF FWHM is not significantly different between the objectives (p = 0.554, Kruskal-Wallis; Fig. 5D, radial).

Soma diameters of mouse pyramidal cells range between 10 to 20 µm, making the cell size, and rather than the optical PSF, the limiting factor regulating the resolution of photostimulation^37^.

Next, we measure the axial PPSF FWHM in the default condition. The 20x, 16x, and 10x objectives yield average axial PPSF FWHMs: 29.3 ± 2.34 µm (‘20x’, N = 43 cells, 4 FOVs, 2 mice), 41.6 ± 6.61 µm (‘16x’, N= 17 cells, 2 FOVs, 2 mice), and 66.4 ± 9.64 µm (‘10x’, N = 12 cells, 2 FOVs, 2 mice) (mean ± SEM; Fig. 5D, axial). The 10x objective yields significantly greater axial PPSF FWHM than the 20x objective (p = 0.0002, Kruskal-Wallis, Bonferroni-corrected). Critically, we see a drop in the 10x axial PPSF FWHM to 39.57 ± 3.27 (N = 32 cells, 8 FOVs, 4 mice) under the swap configuration, making it comparable to the 20x objective under the default configuration (p = 0.12, Kruskal-Wallis, Bonferroni-corrected, non-significant; Fig. 5D).

## Discussion

Here we provide a head-to-head comparison of design choices for multiple 3D-SHOT multiphoton holographic optogenetic systems, so that future users can customize their stimulation systems to fit their needs. We vary the light path, objective, and SLM to evaluate the pros and cons of each decision. We use this approach to design what, to our knowledge, is the largest field of view for *simultaneous* holographic stimulation yet reported, paving the way for new large-scale optogenetic experiments.

While differences in path design, beam width, and SLMs can be modeled, assessing the optical and physiological point-spread function provides a necessary benchmarkfor both custom and commercial multiphoton holographic optogenetic systems. When objectives of varying magnifications are evaluated under identical stimulation and imaging path conditions, they yield significantly different axial profiles, FOV ranges, and power efficiencies. As expected, increasing the objective magnification improves the axial resolution, but minimizes the FOV.

To compensate for this trade-off, we outline a simple, yet effective strategy to maximize the FOV while maintaining hologram axial fidelity. By simply swapping two lenses (L4/L5) in the stimulation optical path, we decrease our axial optical and physiological PSFs at the cost of the accessible stimulation FOV. This allows our ‘10x’ objective to perform similarly to our typical 20x objective, but with a larger imaging FOV.

The need for larger FOVs manifests itself when studying how multiple brain regions interact or theoretically when using larger mammalian species (e.g., rats, ferrets, and non-human primates). As the physical size of the brain scales up exponentially, the size of an individual neuron remains roughly consistent^44^. The ability to scale the FOV while keeping holographic spatial resolution intact allows for optogenetic techniques to manipulate distributed neural circuits across multiple cortical areas simultaneously, without sacrificing single-neuron specificity. Conversely, in smaller animal models with closely packed neurons across a compact FOV (e.g., *Drosophila melanogaster* or *Caenorhabditis elegans*), optimizing the stimulation optical path to achieve finer axial and radial resolution is an essential prerequisite to prevent off-target activation.

The current state of the art in high-resolution, multiphoton holographic photostimulation is structurally bound by a trade-off between optical steering angles and laser power density. In conventional configurations utilizing standard liquid-crystal spatial light modulators (LC-SLMs) and high-NA objectives, the common addressable holographic stimulation within a single FOV is typically restricted to ranges between 200 x 200 µm and 500 x 500 µm^3,4,10,11,13–16,20,29,32,45–53^, with some approaches pushing towards to 1 x 1 mm^1,6–9,19,28,33,39,54,55^. Although higher-density SLMs (e.g., 1920 x 1152 pixels) expand the stimulation range up to 1.3 mm, i.e. outside our standard imaging field of view^41^, it becomes impractical to make use of this whole area. Moreover, while functional imaging with mesoscopes can now capture much larger areas, to date the attempts to use multiphoton holographic optogenetics across a greater area remains restricted to ∼1 x 1 mm albeit moveable within the 5 x 5 mm FOV^40^, or reliant on asynchronous techniques^39^. Consequently, the issue of multi-area, *simultaneous* photostimulation across a large FOV remains.

Recognizing how both custom and commercial rigs can be modified for specific experimental demands highlights the adaptability of these platforms, moving away from a ‘one-size-fits-all’ paradigm toward highly customizable, application-specific configurations. As holographic methods become increasingly prevalent in neuroscience, we demonstrate how a choice as fundamental as objective lens selection directly alters the axial resolution and spatial confinement of the holographic spot.

## Methods

### Two-photon microscope system

*In vivo* two-photon calcium imaging and two-photon holographic optogenetic stimulation was performed using a movable-objective microscope system (MOM, Sutter) combined with a 3D-SHOT system^29,31^. The microscope was equipped with a resonant-galvo scanning system (Sutter Instrument) and a pulsed Ti:sapphire laser (λ = 920 nm, Discovery, Coherent). A motorized three-axis system (MP-285, Sutter) connected with a microscope head enabled locating an objective lens above the targeted brain area. An electrical tunable lens (EL-10-30-TC-NIR-12D, ICC-4C-500, Optotune) was placed in the optical pathway of imaging which enables fast multiplane volumetric imaging. A polarizing beamsplitter (PBS255, Thorlabs) merges the imaging laser and holographic stimulation laser paths. The microscope system was controlled by ScanImage software (MBF Bioscience). Green and red fluorescence was collected with one of four microscope objectives: (1) 20x XLUMPLFLN Objective, 1.0 NA, Olympus, (2) 16x CFI LWD Pan Fluorite, 0.8 NA, Nikon, (3) 10x Super Apochromatic, 0.5 NA, ThorLabs, or (4) 10x Cousa, 0.5 NA, Pacific Optica. Photons were separated by a dichroic mirror (T565lpxr, Chroma) and barrier filters (green: ET525/70m-2p, red: ET605/70m-2p, Chroma) and measured by PMTs (H10770PA-40-04, Hamamatsu photonics). Imaging frames are 512 x 512 pixels. Field of view (FOV) of two-photon images at 1.0x optical zoom were approximately: 1) 1030 × 1030 µm (20x), 2) 1300 x 1300 µm (16x), and 3) 2200 x 2200 µm (10x) for the default configuration. To avoid the excitation of opsin by scanning of the imaging laser (cross-talk)^29,33^, we performed multiplane imaging *in vivo* at a relatively slow frame rate (7.5 Hz) with approximately 50 mW laser power. Imaging power was approximately 15-20 mW when viewing the FOV on a fluorescent slide to avoid bleaching.

### Swap Configuration of the Stimulation Path

In the swapped configuration of the 3D-SHOT stimulation optical path, lens 4 (*f* = 150 mm) and lens 5 (*f* = 100mm) are swapped. As the focal distance between the SLM and the newly replaced lens 4 are now smaller, both lenses are moved closer to the SLM while keeping the focal distance between lens 4 and 5 the same. To move the best resolution holograms to an axial depth between 0 - 100 µm from the primary imaging plane (plane 0), lens 6 is moved to the best possible focal distance between lens 5 and the tube lens, although this change. Only the ‘Olympus – 20x’ objective and the ‘Cousa – 10x’ objectives are tested in this configuration. The HSP1k and HSP1920 were used for data shown in Figure 3 and Figure 4, respectively.

### Calibration procedure

The calibration procedure, adapted from^7,30^, is run using custom MATLAB-scripts, functioning to synchronize the procedure with ScanImage, imaging and stimulation laser emission, and SLM phase mask control. In setting up for a calibration, a thin fluorescent slide is placed atop a cage system housing a 4x objective, an optical lens (*f* = 125 mm), a bandpass filter, and a CMOS camera (CS2100M-USB, ThorLabs). Objectives are then lowered to their respective working distances and aligned with the substage setup to ensure the imaging FOV is visible on the center of the camera.

The calibration pipeline operates sequentially through the following steps: 1) The Sutter steps through defined z-depths while a matrix grid of spots evaluates the fluorescence intensities of the imaging field of view seen on the camera, establishing the coordinate transformation between the raw camera pixels, optical depths, and the focal planes for each ETL plane. A fast coarse z-stack sweep determines localized coordinates and intensity thresholds for 200 individual targets, followed by an optimized multi-target fine search grouping nearby targets into spatially separated, non-overlapping multi-hologram clusters (700 total targets: 20 targets per hologram, 5 distinct sets of targets per imaging plane, 7 planes). Each target was fit with a gaussian to determine the axial and radial FWHM, with the peak representing target fluorescence intensity. A polynomial model defines the transformation between arbitrary ‘SLM coordinates’ to locations in camera space. After calculating the change-in-coordinates (CoC) between camera and SLM space, approximately 50 candidate targets corresponding to each imaging plane are selected. Using a higher laser intensity the fluorescent slide is bleached by each holographic target one at a time. After bleaching holes, images of the fluorescent slide are consecutively registered alongside the expected coordinate, and a second polynomial model is created to relate ‘SLM coordinates’ to locations on 2p images.

### Data Preprocessing and Quality Control

To ensure high-fidelity spatial characterization, calibration measurements were excluded from downstream analysis if: 1) Targets with sensor intensity values exceeding the camera maximum saturation threshold, 2) data points outside the targeted experimental depth (< -25 or >120 µm), and 3) data that was poorly fit by a gaussian.

***Power Efficiency:*** Target power efficiency was determined by normalizing the measured intensity against the wavelength-dependent optical transmission of the objective. To normalize metrics across configurations, intensity values were scaled relative to the 99^th^ percentile across the FOV, with values bounded between 0 and 1.

### Lateral FOV Profile

To map the performance falloff across the lateral FOV, all spatial profiles for power efficiency and FWHM were rendered along X and Y axes (Figures 2 and 4). Sorted X- and Y-lateral distributions were fitted with a gaussian. The lateral center of the FOV was defined by the peak center location of the efficiency fit, and all spatial coordinates were re-centered relative to the origin. Spatial Full Width at 20% Maximum (W_0.2_) was derived, representing the lateral FOV diameter before efficiency drops below 80% of its peak value. To evaluate optical performance across calibration depth (SFig. 1), target metrics (power efficiency, axial/radial FWHM) were separated by depth and FOV radius. Measurements were assigned to three depth bins centered at 0, 50, and 100 µm (edges at -25, 25, 75, and 125 µm). To analyze FOV fall-off, Euclidean radial distances from the FOV center of each objective were calculated and grouped into concentric radial ring zones spaced at 200 um intervals up to 1.2 mm. Spots with power efficiencies below 0.2 (20%) were excluded from characterization.

### Animals

All experiments were performed with approval of the Animal Care and Use Committee of Rutgers University, USA, and in accordance with IACUC guidelines, protocol number #PROTO20220005. TetO-GCaMP6s (Jax stock #024742) mice crossed with Camk2a-tTA mice (Jax stock #003010) were used in this study. Male and female animals at least older than 6 weeks-old at the start of the experiment were used throughout this study. Animals were housed on a reversed 12-hour dark / light cycle at an ambient temperature (22 °C) and humidity (55 %) and had free access to food and water. No statistical methods were applied to pre-determine the sample size of mice for each experiment.

### Surgical procedure

Mice were anesthetized by isoflurane (5% induction, then 1.5-2.0% maintenance) through oxygen-enriched air. Mice were secured in a stereotaxic frame (Model 1920, Kopf) and kept warm with a heating pad (Rodent Warmer X1, Stoelting). Their eyes were covered by a protective eye lubricant during the surgery. Extended-Release Buprenorphine (Ethiqa XR, 3.25 mg/kg) was subcutaneously injected for analgesia, and bupivacaine (3 mg/kg) was injected under the skin over the skull for local anesthesia. Dexamethasone (2 mg/kg) was subcutaneously injected to reduce cerebral edema during craniotomy. A custom-designed titanium headplate was implanted on the skull. The headplate was secured with C&B Metabond (Parkell). A craniotomy was made over the central forelimb area of mouse motor cortex (CFA), coordinates from bregma 0.3 mm anterior, 1.5 mm lateral, 350 µm deep^56^. For the injection of adeno-associated virus (AAV), TetO-GCaMP6s mice crossed Camk2a-tTA mice were injected with AAV php.eb.TRE.stChroME2s-mRuby3 (Virovek, ca 150-300nl, diluted by sterile PBS, 1.3 - 2.5e^12^ vg/ml) into CFA. The glass pipette was kept in place for 2-5 minutes after the AAV injection to ensure adequate spread and prevent backflow of AAV during retraction of the pipette. A cranial window was constructed from two 2.0 mm and one 3.0 mm diameter circular coverslips (Thickness No. 1, Matsunami US). The fabricated coverslip was placed onto the craniotomy and secured into place with Metabond.

### In vivo two-photon holographic optogenetics

*In vivo* holographic two-photon optogenetics was performed using the custom-designed 3D-SHOT system simultaneous with functional two-photon imaging. The details of the default 3D-SHOT system are as described previously^7,28,29^. Briefly, we set up the two-photon holographic optogenetics pathway with a pulsed laser (λ = 1030 nm, 50W, aeroPULSE FS50, NKT Photonics) at low repetition rate (approximately 1.25 MHz) and a fast spatial light modulator (HSP1k-1064-HSP8-TCS2, 1024 × 1024 pixels, Meadowlark Optics; HSP1920-1064-HSP8-HB, 1920 × 1152 pixels; Meadowlark Optics). In each holographic optogenetics session, a phase mask was generated on the SLM using custom-written MATLAB code to make a 3D hologram. The stimulation laser was gated to only illuminate around the imaging system’s galvo mirror flyback period, restricting stimulation artifacts to the edges of the two-photon images^7,29^. With the laser gate, the stimulation artifact was minimized on the lateral edges in the two-photon images.

### Setting a custom power per cell

To compensate for the variable intrinsic excitability and the level of opsin expression across neurons, we determined an individual power to drive each targeted neurons^7,28,29^. Briefly, in each experiment, we stimulated groups of neurons with a train of five 5ms pulses at various intensities. The calcium responses were analyzed online using either custom-written MATLAB codes or an online interpretation of the CaImAn onACID algorithm^57,58^ adapted for MATLAB^7^. The target power used for each cell during the rest of the experiment was set as the minimal power necessary to drive that cell to the saturating level. Neurons were then selected for the physiological point spread function experiments, and those not reliably activated during this procedure were not used.

### Physiological point-spread function measurements

To measure the resolution of our holographic targeting, we measured the physiological point spread function (PPSF) using each objective^7^. We first performed a power test, followed by the random selection of two groups of ten photoactivatable neurons in the FOV. Holograms were offset radially in 3 µm steps (range: -3 to 30 µm) and axially in 6 µm steps (range: -6 to 60 µm) from the target neuron. The resulting calcium activity (ΔF/F) is fit to a gaussian, and the full-width half max (FWHM) was calculated. Error bars are SEM.

### Image processing

Two-photon images were processed using Suite2P^59^. The images were motion-corrected, and regions of interest (ROIs) were automatically generated. Next, experimenters curated ROIs and sorted them as neurons or not. A neuropil signal was calculated for each ROI via Suite2p, and each ROI signal was subtracted from the neuropil signal multiplied by a correction coefficient (0.7). The resulting signals represent the raw fluorescent calcium trace for individual neurons. Raw fluorescence was converted to ΔF/F, using the 10th percentile of a moving baseline to determine F0. The resulting ΔF/F vector was converted to z-ΔF/F via z-scoring of the unrolled fluorescent trace.

### Exclusion Criteria

We excluded the data similarly as described previously^7^. Trials were excluded in case 1) half or more than half of the targeted neurons failed to respond to the holographic stimulation (beyond 0.25 z-ΔF/F); 2) the brain was found to have moved more than 4.8 µm (3 pixels) during stimulation. Neurons were excluded in case 1) they were found in an “off-target” region, 15 µm from targeted neurons radially and 30 µm radially from the targeted neurons from one imaging plane away; 2) non-targeted neurons were located within a stimulation laser artifact area. Neural ensembles were excluded if more than 50% of the targeted neurons were not detected via the suite2p process, or if more than 50% of trials were excluded from that condition. For PPSF experiments, neurons were excluded from the analysis if the resulting gaussian fits failed to achieve a coefficient of determination (R^2^ ≥ 0.5) or yield an extreme peak center estimate (|b_1_| > 100 µm).

### Analytical Model

To systematically evaluate and optimize system performance across different objective lenses and optical configurations, we developed an analytical model of the 3D-SHOT beam path. The model tracks beam expansion and spectral dispersion at each step of the light path, deriving the radial and axial FWHM as well as the maximum addressable FOV for stimulation.

### Statistical analysis

Statistical analysis was performed using MATLAB (MathWorks). Alpha level was set at 0.05. Data are presented as mean ± SEM unless otherwise stated. To test the effect of axial depth and FOV radial zones across objectives while accounting for baseline objective differences (i.e., axial FWHM), data were analyzed using a pooled linear mixed-effects (LME) model. Depth (µm) and radial distance (mm) were modeled as continuous fixed-effect predictors, while individual objectives were included as a random intercept. PPSF group differences were assessed using a non-parametric Kruskal-Wallis test followed by post-hoc pairwise comparisons with Bonferroni-adjusted p-values.

## Supporting information

Supplementary Model

## Data availability

Source data to generate figures will be provided upon publication.

## Code availability

The code used for calibration, analysis, and figure generation is available on Github: OldenburgLab-Public.

## Acknowledgements

We thank Sergey Matveev from ThorLabs for lending us the 10X Apochromatic objective. This work is supported by the CSCR26FEL012 from the NJ Commission on Spinal Cord Research to B.G.; Brain & Behavior Research Foundation Young Investigator Grant to M.H.; funds from the Whitehall Foundation, the Searle Scholars Program, NIH R00EY029758, and DP2MH140135 to I.A.O.

## Supplementary Materials

**Supplementary Table 1.** Objectives’ Reported Optical Properties.

| Objective | Magnification | Numerical Aperture | Effective Focal Length (mm) | Working Distance (mm) | Immersion Medium |
| --- | --- | --- | --- | --- | --- |
| Olympus XLUMPLFN | 20 | 1.0 | 9.0 | 2.0 | Water |
| Nikon CFI LWD Pan Fluorite | 16 | 0.8 | 12.5 | 3.0 | Water |
| ThorLabs Super Apochromatic | 10 | 0.5 | 20 | 7.77 | Air |
| Pacific Optica Cousa | 10 | 0.5 | 20 | 20.0 | Air |

**Supplementary Table 2.** Measured OPSFs versus Modeled OPSFs.

| SLM | Objective | Measured |  |  | Modeled |  |  |
| --- | --- | --- | --- | --- | --- | --- | --- |
|  |  | Radial FWHM* (μm) | Axial FWHM* (μm) | Stim FOV (mm) | Radial FWHM (μm) | Axial FWHM (μm) | Stim FOV (mm) |
| HSP 1K | Olympus - 20x | 9.1 | 17.0 | 0.700 | 8.00 | 10.4 | 0.91 |
|  | Nikon - 16x | 9.3 | 27.5 | 0.976 | 11.1 | 20.3 | 1.26 |
|  | Thorlabs - 10x | 11.5 | 55.8 | 1.81 | 17.8 | 52.3 | 2.02 |
|  | Cousa - 10x | 12.2 | 45.2 | 1.80 | 17.8 | 52.3 | 2.02 |
| HSP 1920 | Olympus - 20x | 9.00 | 16.9 | 0.93 | 8.00 | 10.4 | 1.69 |
|  | Cousa - 10x | 10.0 | 42.8 | 2.24 | 17.8 | 52.3 | 3.75 |
|  | Olympus - 20x (swap) | 8.8 | 9.5 | 0.67 | 3.56 | 1.82 | 0.75 |
|  | Cousa - 10x (swap) | 8.7 | 15.6 | 1.25 | 7.90 | 10.1 | 1.66 |

\*Averaged FWHM values for targets with power efficiencies > 0.2, representing the center of the field of view for a more realistic comparison to modeled values.

**Supplementary Figure 1.**
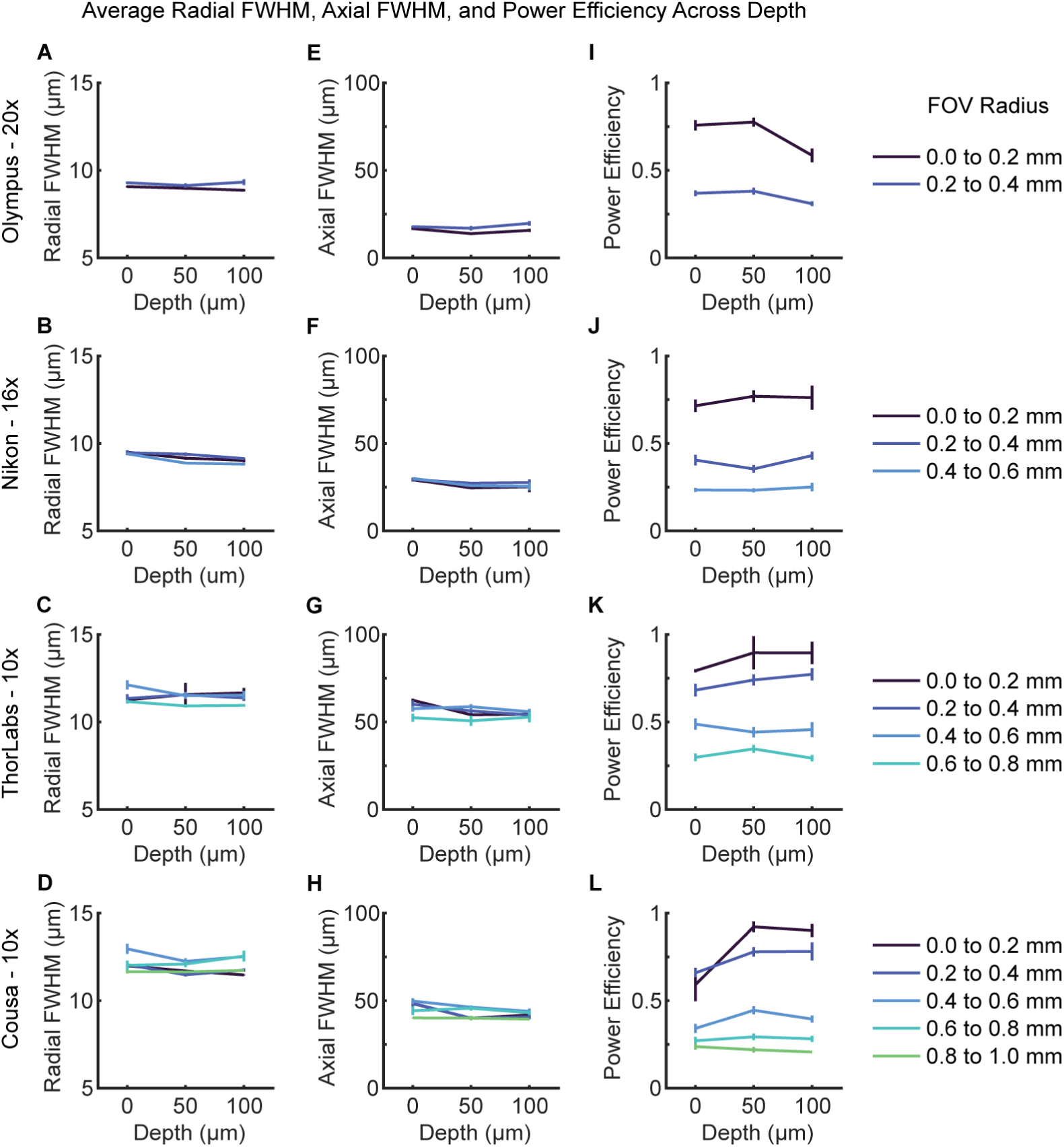
Optical performance across depths and FOV radial zones. **A-D)** Radial FWHM (µm) plotted against depth across concentric radial ring zones (0.2 mm increments, indicated by color) for each objective (rows 1-4). **E-H)** Corresponding axial FWHM (µm) across depth and FOV zones. **I-L)** Hologram power efficiency across depth and FOV zones. Data points represent binned means ± SEM across targets within each FOV zone. Data points with power efficiencies ≤ 20% power efficiency are excluded from the analysis.

**Supplementary Figure 2.**
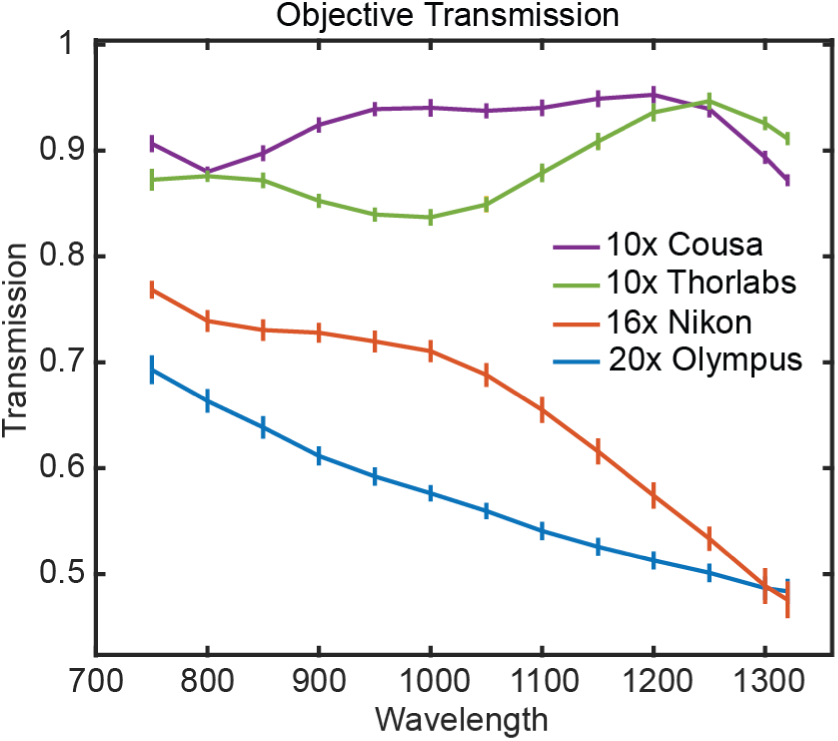
Objective transmission as a function of wavelength (750 – 1320nm) for four objectives: 20x Olympus (blue), 16x Nikon (orange), 10x Cousa (purple), and 10x Thorlabs (green). Transmission was calculated as the ratio of power measured immediately after the objective to the power measured directly before the objective. Data points represent the mean ± SEM across n=5 independent measurement replicates per wavelength.

